# A single-nucleus and spatial transcriptomic atlas of the Syrian hamster preoptic area across hibernation states

**DOI:** 10.64898/2026.09.04.749515

**Authors:** Haruaki Sato, Hikaru Sugimoto, Chengru Shao, Akari Yamauchi, Masafumi Tsurutani, Teppei Goto, Mitsue Hagihara, Dooseon Cho, Naoko Shimada, Mitsutaka Kadota, Takefumi Kondo, Shinya Kuroda, Genshiro A. Sunagawa, Hiroshi Kiyonari, Yoshifumi Yamaguchi, Kazunari Miyamichi

## Abstract

Hibernation is an extreme physiological state in certain mammals that is characterized by a reversible reduction in metabolic and thermogenic demands. Although the preoptic area has been implicated in the regulation of hibernation, comprehensive molecular and spatial resources for this brain region in hibernating mammals remain limited. Here, we present an integrated transcriptomic resource of the preoptic area generated using single-nucleus RNA sequencing (snRNA-seq) and Xenium-based spatial transcriptomics in a mammalian hibernator, Syrian hamster. The resource includes cross-species transcriptomic annotations relative to mouse preoptic area datasets, seasonal snRNA-seq datasets obtained under non-hibernating and hibernating conditions, and spatially resolved cell-type annotations generated from Xenium profiling. Together, these datasets provide a reference framework for identifying neuronal populations, examining molecular conservation across species and conditions, and integrating transcriptomic identity with spatial organization.

## Background & summary

Certain mammalian species living in harsh environments can enter the distinctive physiological state of hibernation, which enables survival for several months in the absence of warmth or food and water^1,2^. Hibernation is defined by active and drastic suppression of metabolism and thermogenesis and often involves prolonged (>24 h) bouts of torpor interspersed with periodic arousals^3,4^. Deep torpor bouts are periods of profound energy conservation characterized by marked reductions in metabolic and thermogenic demands, during which body temperature falls below 10 °C. Hibernation occurs across diverse mammalian lineages^5^, including primates^6^, raising the possibility that hibernation-like states might be inducible in humans. This prospect has attracted considerable attention owing to its potential medical applications^7^. However, the neuronal and physiological mechanisms that enable hibernation remain incompletely understood, posing a major barrier to exploring its translational potential.

Decades of research using hibernating species such as squirrels and hamsters have begun to elucidate the neuronal mechanisms underlying hibernation^3,4^. For example, relatively broad cellular lesions including the preoptic area (POA) disrupt inter-torpor arousals in ground squirrels^8^. In these species, c-Fos, a proxy for neuronal activation, is selectively upregulated in the medial preoptic area during the induction phase of torpor^9^. Collectively, these observations imply that POA neurons may contribute to both the induction and arousal phases of hibernation. In addition, a series of pharmacological studies has demonstrated the importance of several neuromodulatory and neuropeptide signals, including adenosine, opioids, endorphins, and thyrotropin-releasing hormone, many of which are thought to act on POA and hypothalamic neurons^10–13^. However, a major weakness of these studies is their limited spatial and cell-type resolution. Recently, seminal studies have begun to utilize single-cell transcriptome analyses^14^ and high spatiotemporal-resolution *in vivo* Ca^2+^ imaging^15^ in hibernating animals, providing deeper insight into how appetite and thirst are suppressed during hibernation^16^. To further advance a detailed molecular and cellular understanding of the neuronal control of hibernation, the continued development and application of cell-type-specific neuroscience toolkits^17^ that enable visualization of neuronal activity, mapping of neuronal connectivity, and manipulation of cellular function are warranted.

Regarding thermoregulatory mechanisms in the brain, recent molecular genetic studies in mice, non-hibernating heterotherm species that can exhibit torpor, have provided a detailed cell-type-specific view^18,19^. For example, POA neurons expressing pituitary adenylate cyclase-activating polypeptide (PACAP) and brain-derived neurotrophic factor (BDNF) are activated by warm ambient temperatures and can induce hypothermia upon activation^20^. Prostaglandin E receptor type 3 (EP3R)-expressing neurons mediate febrile responses^21^. POA neurons expressing violet light-sensitive opsin 5 (OPN5)^22^, pyroglutamylated RFamide peptide (QRFP)^23^, leptin receptor^24^, transient receptor potential cation channel subfamily M member 2 (TRPM2)^25^, estrogen receptor alpha (ERα)^26^ or those activated during daily torpor^27^ can each induce a prolonged torpor-like hypothermic state with temporal precision. Notably, many of these thermoregulatory neurons co-express vesicular glutamate transporter type 2 (VGLUT2)^28^, consistent with findings that VGLUT2-expressing neurons can also drive a torpor-like state^24,29^. In addition, cold-sensitive POA neurons expressing bombesin-like receptor 3 (BRS3) are involved in promoting thermogenesis^30^ and also play an important role in coordinating appetite and maternal behaviors^31^. Collectively, these studies highlight the power of mouse genetics to dissect the cellular and molecular mechanisms underlying thermoregulation within the POA.

In contrast to rapid advances in the cell-type-resolved understanding of thermoregulatory systems in mice, a substantial knowledge gap remains in hibernating species such as squirrels and hamsters. It is unclear whether the molecularly defined thermoregulatory cell types identified in mice are conserved in hibernators, or whether hibernators have evolved specialized neuronal populations that enable the extreme reductions in body temperature as observed in deep torpor. Addressing this question is a prerequisite for introducing molecular genetic approaches to dissect hibernation-related neuronal populations in the POA of hibernators.

To address this issue, we first applied single cell transcriptomics^32,33^, specifically single-nucleus RNA sequencing (snRNA-seq), to the POA of the Syrian hamster, a well-established hibernation model^1,2^. We chose the hamster because our collaborative team has recently developed genome-editing techniques in hamster oocytes that enable the generation of Cre recombinase knock-in hamsters^34^. This advance will accelerate the implementation of viral genetic tools^17^ to visualize and manipulate genetically defined neurons in the hamster brain. By generating snRNA-seq data from the hamster POA and comparing them with datasets previously characterized in mice^28,35^, we sought to establish a cell-type census of hamster thermoregulatory neurons. In addition, we compared the transcriptomic profiles of hamsters housed under warm conditions (warm ambient temperature and long photoperiod) with those of hibernating animals in winter-like conditions (cold ambient temperature and short photoperiod), thereby identifying candidate molecular signatures associated with the hibernating state. Finally, we generated a spatial atlas of the hamster POA utilizing 10x Xenium-based spatial transcriptomics^36^ (STx), integrated with snRNA-seq cell-type profiling, to map the spatial organization of thermoregulatory neurons. Together, these datasets provide a reference resource for investigating the molecular identities, transcriptomic conservation, and spatial organization of thermoregulatory neuronal populations in hibernating hamsters.

## Methods

### Animals

All animal experiments were approved by the Institutional Animal Care and Use Committees of the RIKEN Kobe Campus and Hokkaido University. Male and female Syrian hamsters used for snRNA-seq were purchased from Japan SLC, Inc. (Shizuoka, Japan). Non-hibernating hamsters used for snRNA-seq were maintained at ambient temperature (21–24 °C) under a 12-h light/12-h dark cycle before sample preparation. Hibernating hamsters used for snRNA-seq were maintained under cold conditions (4 °C) with an 8-h light/16-h dark cycle at the animal facility of the RIKEN Center for Biosystems Dynamics Research. Hamsters maintained under summer-like conditions (14-h light/10-h dark at 25 °C) and hibernating hamsters (8-h light/16-h dark at 4 °C) used for STx were maintained as described previously^37^ at Hokkaido University. Unless otherwise stated, hamsters were given ad libitum access to a laboratory diet (MFG; Oriental Yeast, Shiga, Japan; 3.57 kcal/g, or MR standard; Nihon Nosan, Kanagawa, Japan; 3.45 kcal/g) and water.

### Single-nucleus RNA sequencing (snRNA-seq): library preparation

The “Frankenstein” protocol (doi:10.17504/protocols.io.3fkgjkw) was modified for nuclei isolation as follows. For the data shown in Figs. 1–3, six male Syrian hamsters at 10 weeks of age were deeply anesthetized with isoflurane (Fujifilm, #099-06571), perfused with cold phosphate-buffered saline (PBS) to remove blood cells, and euthanized by decapitation. Brains were sectioned into 1-mm coronal slices under a microscope, and the sections were floated in 1% BSA-PBS (Nacalai Tesque, #0128197; Takara, #T9181). The POA was dissected based on the location of the anterior commissure. For pooled samples, dissected POA tissues were combined prior to homogenization and thoroughly homogenized using the Minute Single Nucleus Isolation Kit for Tissues/Cells (Invent Biotechnologies Inc., SN-047). Isolated nuclei were resuspended in 1% BSA-PBS containing 0.2 U/μL RNase inhibitor (Roche, #3335399001) and 10 μg/mL 4’,6-diamidino-2-phenylindole (DAPI), incubated on ice for 5 min, and filtered through a 40-μm strainer. Nuclei were sorted from the suspension using a cell sorter (SH800Z; Sony) at 5 °C. Gates were first set to identify single nuclei based on the DAPI signal and then to exclude smaller nuclei, inferred to be glial nuclei, as well as multiplets. Immediately after sorting, nuclei were centrifuged at 600 × g for 5 min at 4 °C. The number of nuclei was then counted using a hemocytometer to verify the yield of intact nuclei. Nuclei were resuspended in PBS and used to generate snRNA-seq libraries using the 10x Genomics Chromium platform, targeting 10,000 nuclei per sample. Libraries were prepared using the 10x Genomics Chromium Single Cell 3′ Reagent Kits v3 (PN-1000269). Completed libraries were sequenced to a depth of 100 Gb on an Illumina NovaSeq 6000 platform by Azenta.

**Figure 1.**
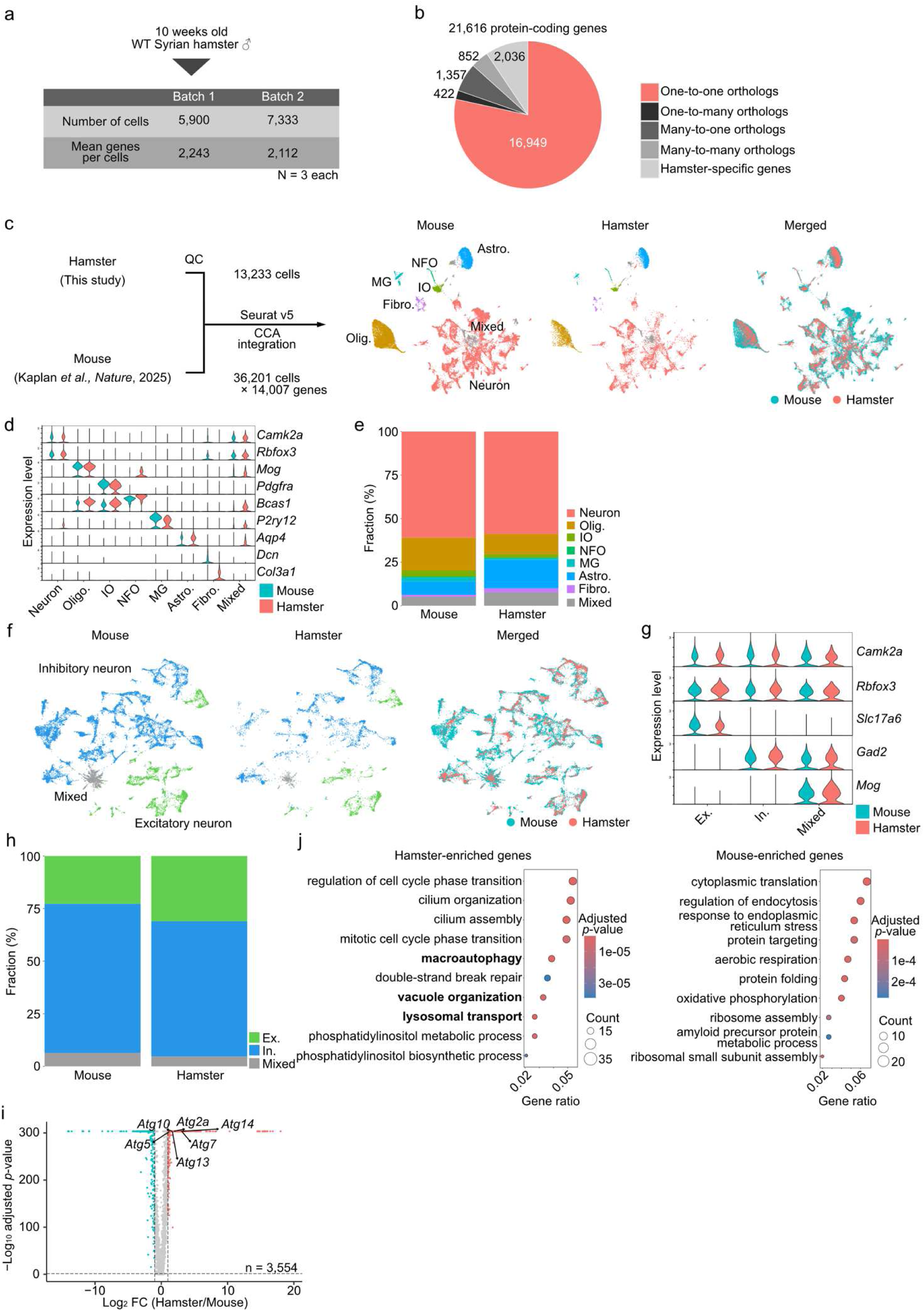
Shared cellular architecture of the POA in hamsters and mice. (**a**) Schematic summary of snRNA-seq metrics in hamsters after quality control. (**b**) Fraction of mapping patterns of protein-coding genes in hamsters identified by OrthoFinder2, compared with mice. (**c**) Schematic overview of the comparative transcriptomics analysis between mice and hamsters (left), and Uniform Manifold Approximation and Projection (UMAP) representation of all cells from hamsters, mice, and the merged dataset (right). Colors in the hamster and mouse UMAPs indicate cell classes, whereas those in the merged UMAP indicate species. Abbreviations: Oligo, oligodendrocytes; IO, immature oligodendrocytes; NFO, newly formed oligodendrocytes; astro, astrocytes; MG, microglia. (**d**) Violin plots showing the expression levels of marker genes for major cell classes in both species. (**e**) Fraction of each cell class in each species. (**f**) UMAP representation of neuronal populations in hamsters (left), mice (middle), and the merged dataset (right). Colors in the left and middle panels indicate excitatory and inhibitory neurons, whereas those in the right panel indicate species. (**g**) Violin plots showing the expression levels of marker genes for excitatory and inhibitory neurons, including excitatory, inhibitory, and mixed subtypes. (**h**) Fraction of excitatory and inhibitory neurons in each species. (**i**) Volcano plots showing species-DEGs enriched in each species across all cells. Selected autophagy-related genes are indicated. The x-axis shows log_2_-transformed fold change, and the y-axis shows −log_10_-transformed *p*-values from the Wilcoxon rank-sum test with Bonferroni correction. Genes with a fold change > 2 or < 0.5 and *p* < 0.05 were defined as species-DEGs. Red and blue dots indicate hamster- and mouse-enriched species-DEGs, respectively. (**j**) Heatmaps of *p*-values from GO analysis (biological process) based on hamster-enriched (left) and mouse-enriched (right) species-DEGs across all cells. Enrichment was assessed using a hypergeometric test, and *p*-values were adjusted for multiple testing using the Benjamini–Hochberg (BH) method.

**Figure 2.**
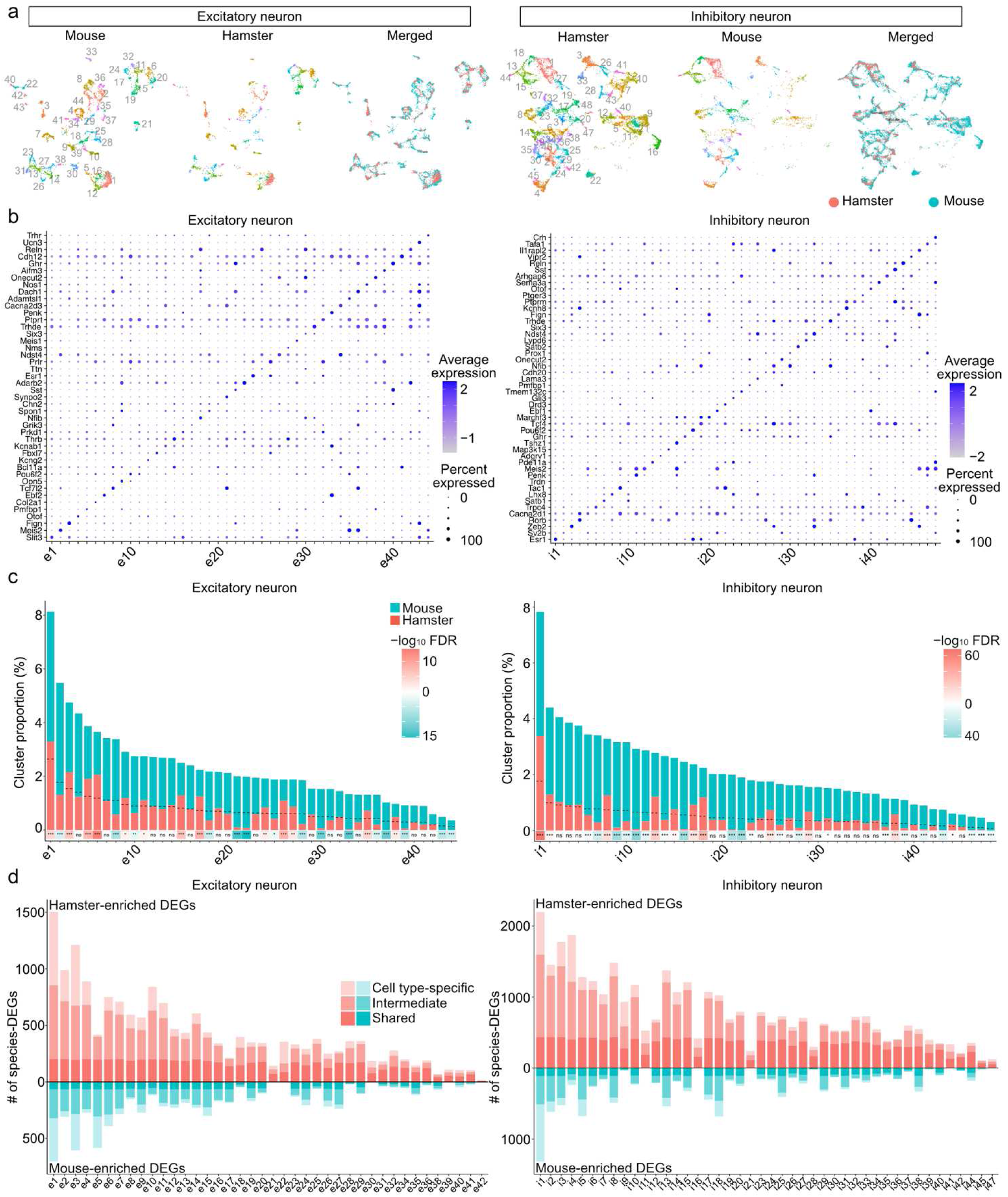
Cross-species comparison of excitatory and inhibitory neuronal subtypes. (**a**) UMAP representations of excitatory (left) and inhibitory (right) neurons from hamsters, mice, and the integrated dataset. Colors in the hamster and mouse panels indicate neuronal subtypes, whereas colors in the integrated UMAP indicate species. (**b**) Dot plots of z-scored expression levels of marker genes across excitatory (left) and inhibitory (right) neuronal subtypes. (**c**) Bar graphs showing species differences in the relative abundance of each excitatory (left) and inhibitory (right) subtype. Heatmaps below the graphs indicate hamster- and mouse-enriched subtypes in reddish and bluish colors, respectively. The *p*-values were calculated using Fisher’s exact test with Benjamini–Hochberg correction. ns: not significant, \**p* < 0.05, \*\**p* < 0.01, and \*\*\**p* < 0.001. (**d**) Bar graphs showing the number of species-associated differentially expressed genes (species-DEGs) in each subtype. Hamster- and mouse-enriched species-DEGs are shown in reddish and bluish colors, respectively. Species-DEGs detected in more than 50% of all subtypes within each neuronal population were classified as “shared,” those detected in fewer than 10% were classified as “cell type-specific,” and the remainder were classified as “intermediate.” Some subtypes were excluded due to low number of cells in hamster or mouse.

**Figure 3.**
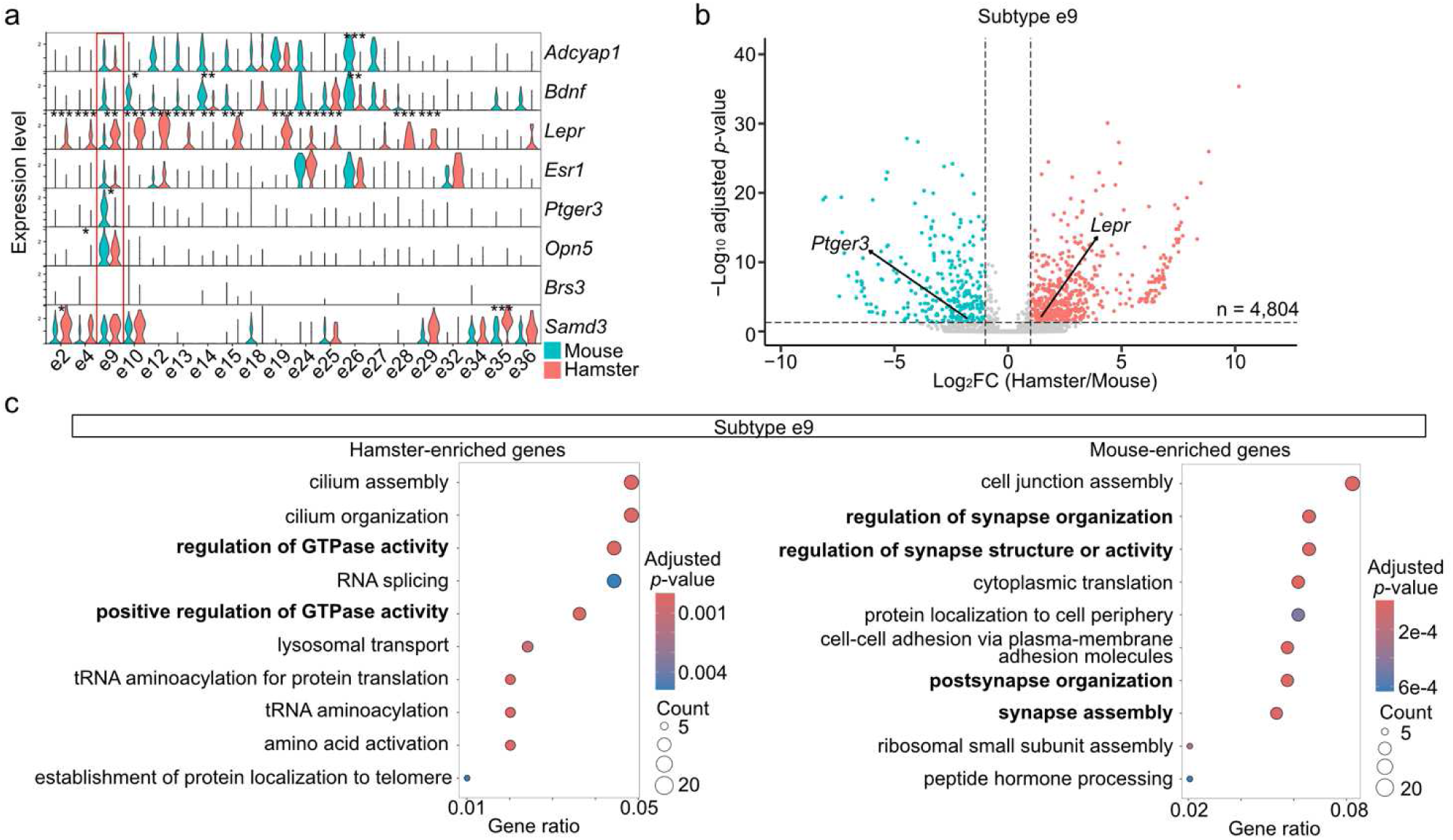
Cross-species comparison of thermoregulatory neuronal subtypes. (**a**) Violin plots showing expression levels of marker genes associated with thermoregulatory neuronal subtypes in the POA of mice. \**p* < 0.05, \*\**p* < 0.01, and \*\*\**p* < 0.001 by the Wilcoxon rank-sum test with Bonferroni correction. (**b**) Volcano plots showing species-DEGs within excitatory*^Opn5^*^/*Ptger3*^ neurons (e9 subtype). The x-axis shows log_2_-transformed fold change, and the y-axis shows −log_10_-transformed *p*-values from the Wilcoxon rank-sum test with Bonferroni correction. Genes with a fold change > 2 or < 0.5 and *p* < 0.05 were defined as differentially expressed genes. Red and blue dots indicate hamster- and mouse-enriched species DEGs, respectively. (**c**) Heatmaps of *p*-values from GO analysis (biological process) based on hamster-enriched (left) and mouse-enriched (right) species-DEGs within excitatory*^Opn5^*^/*Ptger3*^ neurons (e9 subtype). Enrichment was assessed using a hypergeometric test, and *p*-values were adjusted for multiple testing using the Benjamini–Hochberg (BH) method.

For the data shown in Figs. 4–6, non-hibernating and hibernating male and female Syrian hamsters aged 4–8 months were used. Library construction followed the protocol described above, except that the 10x Genomics Chromium Single Cell 3′ v4 GEM-X kit (PN-1000686) was used, with 20,000 nuclei targeted per sample. Completed libraries were sequenced to a depth of 100 Gb on either an Illumina NovaSeq 6000 platform by Azenta or a NovaSeq X Plus platform by A.D.A.M. Innovations; see also Fig. 4a.

**Figure 4.**
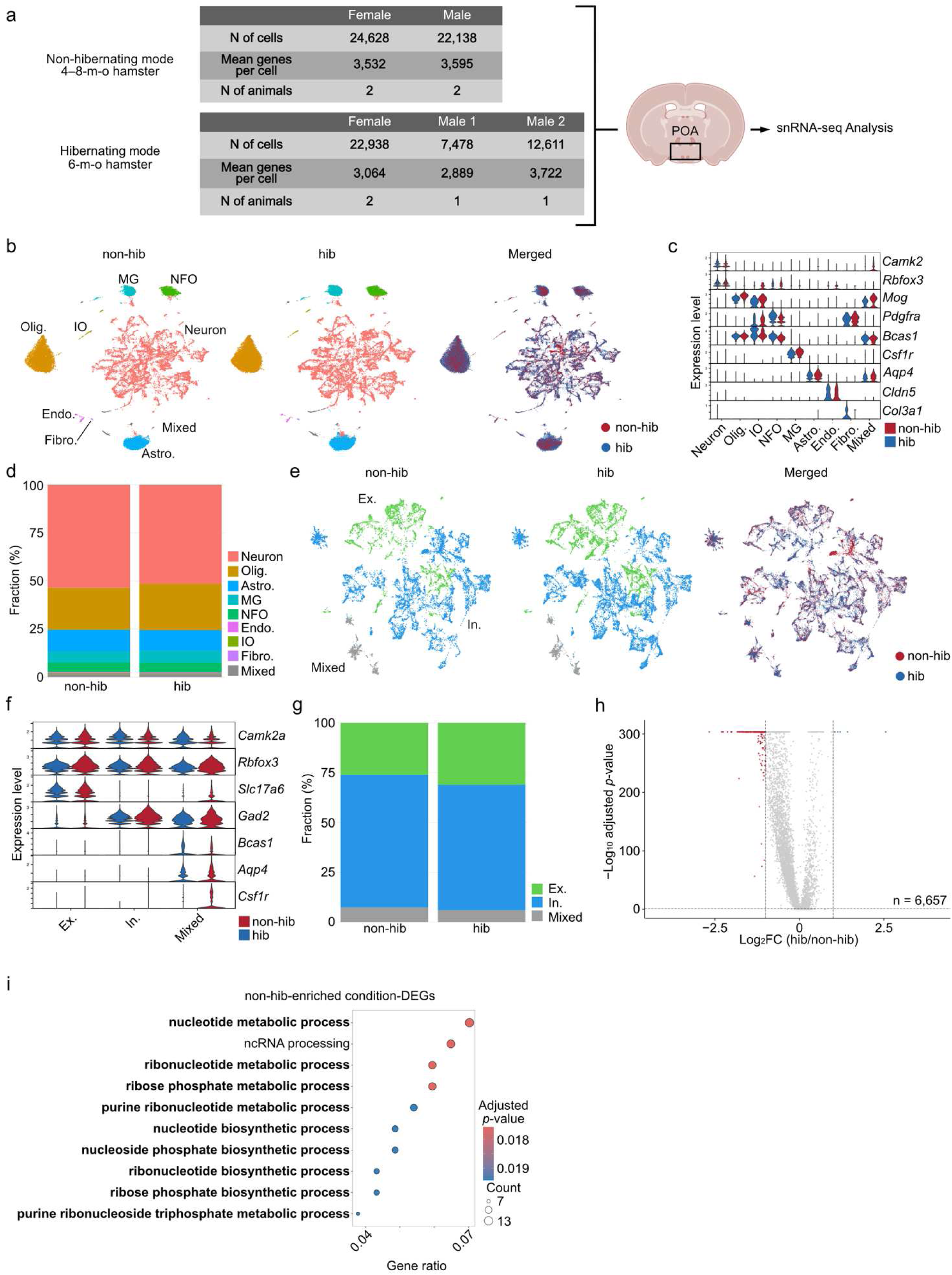
Global impacts of hibernation on POA transcriptomic signatures in hamsters. (**a**) Schematic overview of the snRNA-seq datasets under warm (non-hibernating, non-hib) and winter-like (hibernating, hib) conditions. The table on the right summarizes the dataset after quality control. (**b**) UMAP representation of all cells from non-hibernating hamsters (left), hibernating hamsters (middle), and the merged dataset (right). Colors in the left and middle panels indicate cell classes, whereas those in the right panel indicate physiological state. For abbreviations, see Fig. 1c. (**c**) Violin plots showing the expression levels of cell class marker genes across conditions. (**d**) Fraction of each cell class among total cells. (**e**) UMAP representation of neuronal populations in non-hibernating hamsters (left), hibernating hamsters (middle), and the merged dataset (right). Colors in the left and middle panels indicate excitatory and inhibitory neurons, whereas those in the right panel indicate hibernation state. (**f**) Violin plots showing expression levels of marker genes for excitatory and inhibitory neurons, including excitatory, inhibitory, and mixed subtypes. (**g**) Fraction of excitatory and inhibitory neurons among total neurons. (**h**) Volcano plots showing condition-DEGs downregulated (left) and upregulated (right) during hibernation across total neurons. The x-axis shows log_2_-transformed fold change, and the y-axis shows −log_10_-transformed *p*-values from the Wilcoxon rank-sum test with Bonferroni correction. Genes with a fold change > 2 or < 0.5 and *p* < 0.05 were defined as condition-DEGs. Blue and red dots indicate upregulated and downregulated condition-DEGs during hibernation, respectively. (**i**) Heatmap of *p*-values from GO analysis (biological process) based on condition-DEGs during hibernation in total neurons. Enrichment was assessed using a hypergeometric test, and *p*-values were adjusted for multiple testing using the Benjamini–Hochberg (BH) method.

**Figure 5.**
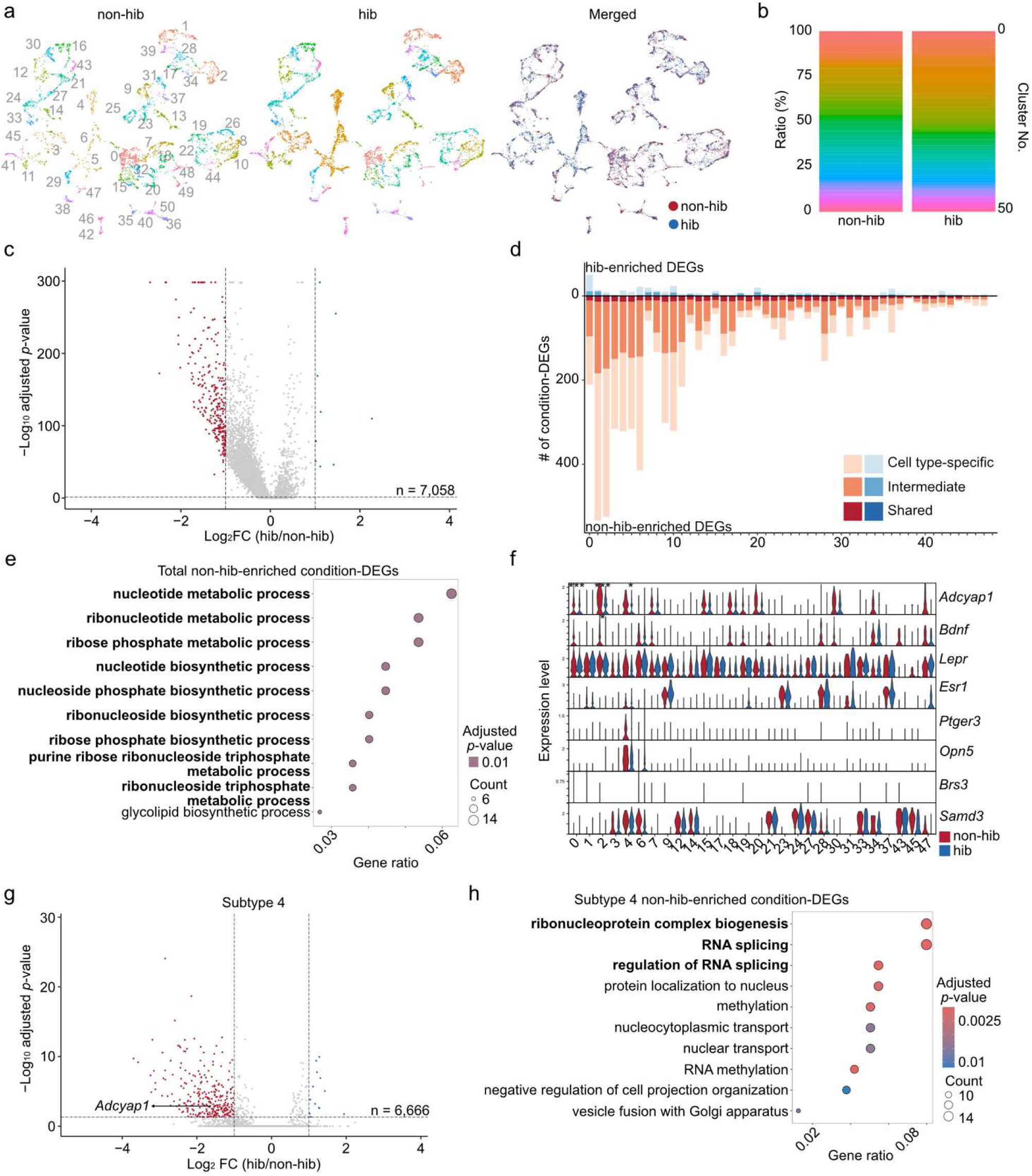
Impact of hibernation on excitatory neuronal subtypes in the POA. (**a**) UMAP representation of excitatory neurons in non-hibernating hamsters (non-hib, left), hibernating hamsters (hib, middle), and the merged dataset (right). Colors in the left and middle panels indicate neuronal subtypes, whereas colors in the right panel indicate hibernation state. (**b**) Bar graphs showing the proportion of each excitatory subtype within the excitatory neuronal population. (**c**) Volcano plots showing condition-DEGs identified in pan-excitatory neurons. The x-axis shows log_2_-transformed fold change, and the y-axis shows −log_10_-transformed *p*-values from the Wilcoxon rank-sum test with Bonferroni correction. Genes with a fold change >2 or <0.5 and *p* < 0.05 were defined as condition-DEGs. Blue and red dots indicate upregulated and downregulated condition-DEGs during hibernation, respectively. (**d**) Bar graph showing the number of condition-DEGs in each excitatory subtype. Bluish and reddish colors indicate upregulated and downregulated condition-DEGs during hibernation, respectively. Categories are defined as in Fig. 2d. (**e**) Heatmaps of *p-*values from GO analysis, biological process, based on downregulated condition-DEGs in pan-excitatory neurons. Enrichment was assessed using a hypergeometric test, and *p*-values were adjusted for multiple testing using the Benjamini–Hochberg (BH) method. (**f**) Violin plots showing expression levels of marker genes previously characterized in mice in the context of POA thermoregulation. \**p* < 0.05, and \*\*\**p* < 0.001 by the Wilcoxon rank-sum test with Bonferroni correction. (**g**) Volcano plots showing condition-DEGs identified in excitatory subtype 4, corresponding to excitatory*^Opn5^*^/*Ptger3*^ neurons. The x-axis shows log_2_-transformed fold change, and the y-axis shows −log_10_-transformed *p*-values from the Wilcoxon rank-sum test with Bonferroni correction. Genes with a fold change > 2 or < 0.5 and *p* < 0.05 were defined as condition-DEGs. Red and blue dots indicate hib- and non-hib-enriched condition-DEGs, respectively. Selected gene name is indicated. (**h**) Heatmap of *p*-values from GO analysis, biological process, based on downregulated condition-DEGs in excitatory subtype 4. Enrichment was assessed using a hypergeometric test, and *p*-values were adjusted for multiple testing using the Benjamini– Hochberg (BH) method.

**Figure 6.**
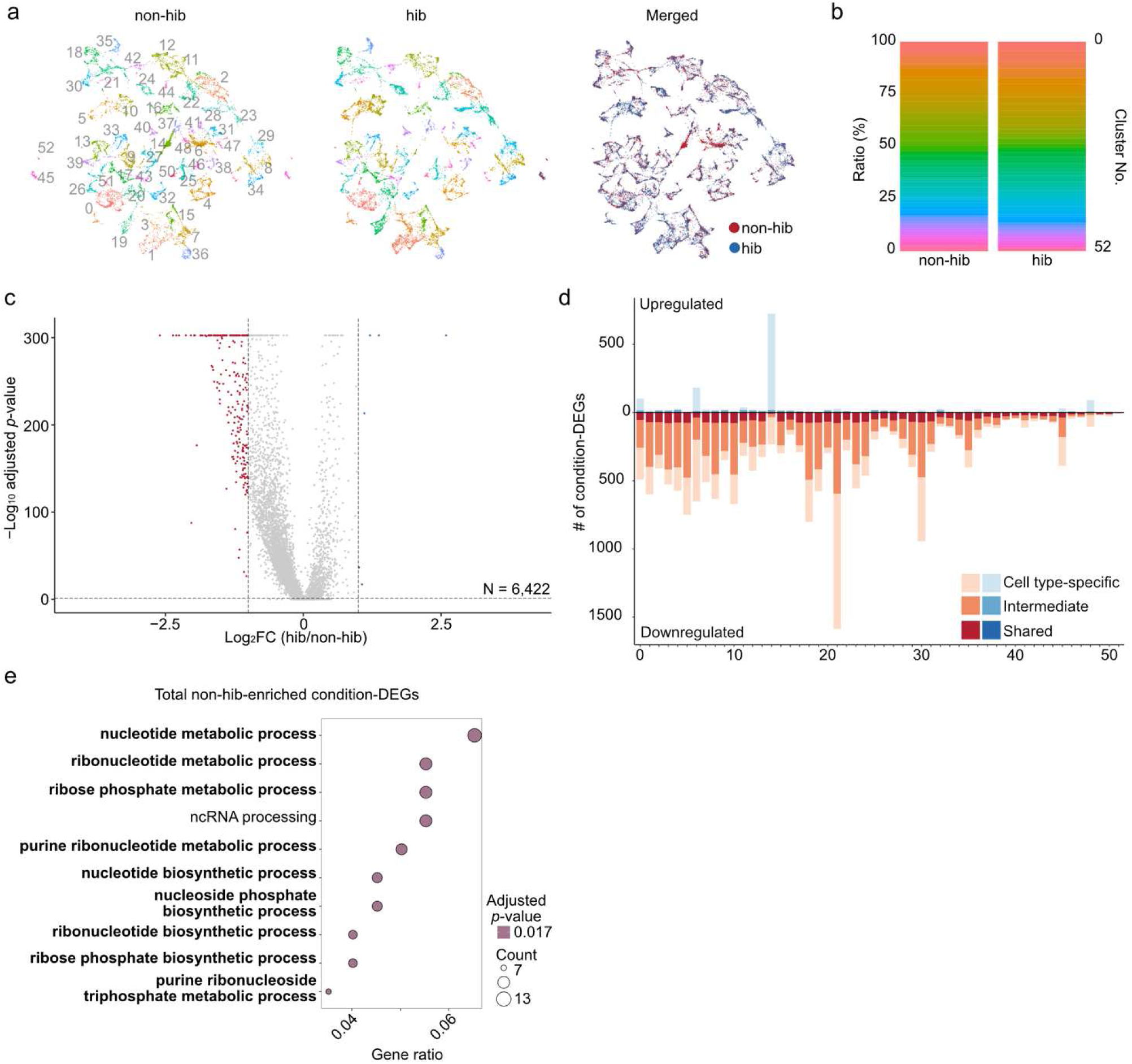
Impact of hibernation on inhibitory neuronal subtypes in the POA. (**a**) UMAP representation of inhibitory neurons in non-hibernating hamsters (non-hib, left), hibernating hamsters (hib, middle), and the merged dataset (right). Colors in the left and middle panels indicate neuronal subtypes, whereas colors in the right panel indicate hibernation state. (**b**) Bar graphs showing the proportion of each inhibitory subtype within the inhibitory neuronal population. (**c**) Volcano plots showing condition-DEGs identified in pan-inhibitory neurons. The x-axis shows log_2_-transformed fold change, and the y-axis shows −log_10_-transformed *p*-values from the Wilcoxon rank-sum test with Bonferroni correction. Genes with a fold change > 2 or < 0.5 and *p* < 0.05 were defined as condition-DEGs. Blue and red dots indicate upregulated and downregulated condition-DEGs during hibernation, respectively. (**d**) Bar graph showing the number of condition-DEGs in each inhibitory subtype. Bluish and reddish colors indicate upregulated and downregulated condition-DEGs during hibernation, respectively. Categories are defined as in Fig. 2d. (**e**) Heatmap of *p*-values from GO analysis, biological process, based on downregulated condition-DEGs in pan-inhibitory neurons. Enrichment was assessed using a hypergeometric test, and *p*-values were adjusted for multiple testing using the Benjamini–Hochberg (BH) method.

### snRNA-seq: data analysis

FASTQ files from each library were first aligned to the reference transcriptome (BCM_Maur_2.0; PRJNA705675) using the Cell Ranger pipeline. After alignment, feature-barcode matrices were loaded into Seurat (R package, v5.3.0)^38^. Nuclei were retained if they contained 400–10,000 unique molecular identifiers (UMIs), with less than 10% of transcript counts from mitochondrial genes and less than 10% from ribosomal genes. Genes were retained if at least one UMI was detected in at least three cells. For cross-species comparison, orthologous relationships were inferred using reference proteomes from 15 mammalian species: golden hamster (*Mesocricetus auratus*, BCM_Maur_2.0), human (*Homo sapiens*, GRCh38.p14), mouse (*Mus musculus*, GRCm39), rat (*Rattus norvegicus*, GRCr8), dog (*Canis lupus familiaris*, UU_Cfam_GSD_1.0), African elephant (*Loxodonta africana*, mLoxAfr1.hap2), lesser hedgehog tenrec (*Echinops telfairi*, ASM31398v2), thirteen-lined ground squirrel (*Ictidomys tridecemlineatus*, HiC_Itri_2), gray mouse lemur (*Microcebus murinus*, Mmur_3.0), little brown bat (*Myotis lucifugus*, Myoluc2.0), squirrel monkey (*Saimiri boliviensis boliviensis*, BCM_Sbol_2.1), Tasmanian devil (*Sarcophilus harrisii*, mSarHar1.11), pig (*Sus scrofa*, Sscrofa11.1), Arctic ground squirrel (*Urocitellus parryii*, ASM342692v1), and American black bear (*Ursus americanus*, gsc_jax_bbear_1.0). Annotated protein sequences corresponding to these assemblies were obtained from NCBI. Orthogroups and orthologs were inferred with OrthoFinder2^39^ (default settings) using DIAMOND^40^ for sequence similarity searches and DendroBLAST^41^ for gene-tree construction. Hamster–mouse gene pairs classified as one-to-one orthologs were extracted and used for downstream cross-species transcriptomic integration. The resulting one-to-one ortholog table is provided in the deposited dataset. snRNA-seq data from mice (GSE280964)^35^ were integrated with the hamster dataset using the Seurat v5 canonical correlation analysis (CCA) integration method after converting hamster gene symbols to the corresponding mouse ortholog names based on the deposited one-to-one ortholog table. The remaining nuclei were normalized using the *LogNormalize* function for cross-species comparison and the *SCTransform* function for cross-condition comparison, with mitochondrial and ribosomal gene ratios regressed out. Dimensionality reduction and integration were performed using *RunPCA*, *IntegrateLayers* with CCA-based integration, *RunUMAP*, *FindNeighbors*, and *FindClusters*, using the first 150 principal components (PCs) and a clustering resolution of 0.4 for cross-species comparison and 0.3 for cross-condition comparison.

Clusters were annotated based on established marker genes for major cell classes, including neurons (*Camk2a*, *Rbfox3*), mature oligodendrocytes (*Mog*), oligodendrocyte precursor cells (*Pdgfra*), newly formed oligodendrocytes (*Bcas1*), endothelial cells (*Cldn5*), fibroblasts (*Col3a1*, *Dcn*), microglia (*P2ry12*), and astrocytes (*Aqp4*). Clusters expressing markers of multiple cell types or containing a low number of detected genes were excluded from further analysis.

For subclustering, neurons were reanalyzed independently. Clustering was performed using 50 PCs with a resolution of 0.4 for cross-species comparison and 0.2 for cross-condition comparison. Doublets and low-quality nuclei were removed using the criteria described above. Neuronal clusters were further classified into excitatory, inhibitory, and cholinergic neurons based on the expression of *Slc17a6*, *Gad1/Gad2*, and cholinergic marker genes, respectively.

Excitatory and inhibitory neurons were then subclustered using 150 PCs. For cross-species comparison, clustering resolutions of 5.5 and 4 were used for excitatory and inhibitory neurons, respectively. For cross-condition comparison, clustering resolutions of 1.8 and 1.6 were used for excitatory and inhibitory neurons, respectively. The clustering resolution was incrementally increased until no additional marker genes capable of distinguishing newly separated clusters could be identified using the *FindMarkers* function. Clusters characterized by similar marker genes were manually merged.

DEGs were identified using the *FindMarkers* function, with thresholds of fold change >2 or <0.5 and P < 0.05. GO analyses were performed using the *clusterProfiler* package, and volcano plots were generated using the *EnhancedVolcano* package in R: https://bioconductor.org/packages/release/bioc/html/EnhancedVolcano.html. The correspondence between cell IDs and annotated cell types or subtypes is provided in the deposited dataset.

### Selection of Xenium custom probes

For selection of Xenium custom probe sets, we used the snRNA-seq dataset generated for cross-species comparison (Figs. 1–3). Marker genes were selected for each excitatory or inhibitory neuronal subtype, excluding clusters that represented less than 0.5% of the population or were more than three-fold more abundant in mice. In addition, major immediate early genes (IEGs; *c-Fos, Arc, Egr1, Homer1, Fosb, Nr4a1, Junb,* and *Dusp1*) and marker genes for major cell classes were included: astrocytes (*Aqp4*, *Slc39a12*), endothelial cells (*Cldn5*), fibroblasts (*Col3a1*), neurons (*Camk2a*, *Rbfox3*), excitatory neurons (*Slc17a6*, *Slc17a8*), inhibitory neurons (*Slc32a1*), oligodendrocytes (*Opalin*, *Plp1*), oligodendrocyte precursor cells (*Pdgfra*), newly formed oligodendrocytes (*Bcas1*), and microglia (*Csf1r*, *Laptm5*). The final set of genes selected for Xenium custom probe design is provided in the deposited dataset.

### Preparations of hamster brain samples for Xenium assay

Three-week-old female Syrian hamsters were purchased from an outbred colony (Japan SLC, Inc., Shizuoka, Japan) and housed under summer-like long-photoperiod and warm conditions (25 °C with a 14 h light/10 h dark cycle), with ad libitum access to water and food (MR standard diet, Nihon Nosan, Japan) as previously described^42^. Three to four individuals were housed per cage, and their body mass was recorded weekly when the cages were replaced. At 10 weeks of age, all animals were transferred to individual polypropylene cages with ad libitum access to water and food. The control group was continuously housed under summer-like conditions thereafter. A Tc logger (iButton, #DS1925L-F5, Maxim Integrated, USA), calibrated by the manufacturer, was coated with rubber (total mass approximately 3.5 g; Plasti Dip, Performix Coatings, USA) and surgically implanted into the abdominal cavity under 2% isoflurane anesthesia before hibernation induction. The logger recorded Tc every 10 min with a resolution of 0.0625 °C. After surgery, animals were allowed to recover for 2 weeks. Four animals were used for each condition. All experiments were performed using adult female hamsters at approximately 7 months of age.

For hibernation induction, the 12-week-old animals were transferred to winter-like short-photoperiod and cold conditions (4–5 °C with an 8 h light/16 h dark cycle), with ad libitum access to water and food. Body mass was measured weekly, and cages were replaced every other week, except for animals undergoing hibernation. Two to three months after transfer to winter-like conditions, the animals began to hibernate and displayed typical torpor–arousal cycles.

To collect brain samples during entry into torpor, 7-month-old hibernating hamsters were placed in cages in which oxygen consumption was monitored in real time using an ARCO-2000 respiratory gas analyzer (ARCO System, Japan). Two hours after a marked reduction in oxygen consumption was observed, the animals were immediately perfused with PBS followed by 4% paraformaldehyde (PFA) in PBS under 2% isoflurane anesthesia. After perfusion, Tc records were retrieved from the iButton, and the brains were collected and post-fixed in 4% PFA in PBS for 1 day at 4 °C. Brains from the control group were collected in parallel and processed in the same manner as those from the hibernating animals. The POA was dissected and subsequently dehydrated in 70% ethanol at 4 °C for at least 1 week. Dehydration and paraffin infiltration were performed using a Tissue-Tek VIP 5 Jr. processor (Sakura Seiki) with the following settings: 90 min in 80% ethanol; 90 min in 90% ethanol; three rounds of 90 min in 100% ethanol; three rounds of 90 min in 50% G-Nox/ethanol (Genostaff, GN08); three rounds of 90 min in G-Nox; and four rounds of 90 min in paraffin (Leica, 39601006). Formalin-fixed, paraffin-embedded (FFPE) blocks were then prepared.

FFPE blocks were sectioned at 5-μm thickness and mounted onto slides according to the manufacturer’s protocol (10x Genomics, CG000578). Sections from all experimental conditions, including all four animals per condition, were placed on each slide using the anterior commissure as a landmark. The number of sections analyzed for each condition and animal is detailed in the deposited dataset.

Xenium analysis was performed using a standalone custom panel (100 genes; 10x Genomics, PN-1000651) and reagents from 10x Genomics (PN-1000460, 1000461, 1000487, and 1000662), following the manufacturer’s protocols (CG000580, CG000749, and CG000584).

### Data analysis of Xenium assay

Output data from Xenium Explorer were processed using Xenium Ranger v3.2. The resulting datasets were imported into R v4.3 and analyzed using the Seurat v5 package. Data were normalized using the *SCTransform* function. Dimensionality reduction and clustering were performed using *RunPCA*, *RunUMAP*, *FindNeighbors*, and *FindClusters*, with 50 PCs and a resolution of 0.3.

Clusters were annotated based on canonical marker genes for neurons (*Camk2a*, *Rbfox3*), excitatory neurons (*Slc17a6*), inhibitory neurons (*Slc32a1*, *Gad1*), oligodendrocytes (*Opalin*), oligodendrocyte precursor cells (*Pdgfra*), microglia (*Csf1r*), fibroblasts (*Col3a1*), endothelial cells (*Cldn5*), ependymal cells (*Adgrv1*), and astrocytes (*Aqp4*). Clusters expressing markers for more than one major cell type were excluded from downstream analyses.

For subclustering of excitatory and inhibitory neurons, each neuronal population was reanalyzed independently using 50 PCs and clustering resolutions of 0.6 and 0.9, respectively. Putative doublets were removed using the same criteria described above. The clustering resolution was incrementally increased until no additional marker genes capable of distinguishing newly separated clusters were detected.

Marker genes for each cluster were identified using the *FindMarkers* function with the following thresholds: adjusted *p* < 0.05, min.pct > 0.1, and log_2_ fold change > 0.25. The correspondence between cell IDs and annotated cell types or subtypes is provided in the deposited dataset. For integration of STx data with the snRNA-seq dataset, RCTD was applied following the Seurat v5 spatial integration workflow (https://satijalab.org/seurat/articles/seurat5_spatial_vignette_2). Pseudo-bulk expression levels for individual animals were calculated using the *AggregateExpression* function in Seurat.

To assess *c-Fos* expression, neurons with *c-Fos* expression levels greater than two standard deviations above the mean across all neurons were defined as *c-Fos*+. The IEG score was calculated using the *AddModuleScore* function in Seurat, and neurons with IEG scores greater than two standard deviations above the mean across all neurons were classified as IEG+. DEG identification and GO analyses were performed in the same manner as described for the snRNA-seq analysis.

## Data Records

Raw paired-end sequencing reads and gene-expression matrices from all samples used for the cross-species and cross-condition comparisons were deposited in the NCBI Gene Expression Omnibus (GEO) under accession number GSE345405 and GSE346081, respectively.

Each gene-expression matrix comprises three compressed text files: barcodes, features (genes), and the sparse count matrix. Additionally, raw data for STx have been stored in the NCBI GEO under accession number GSE345359.

## Technical Validation

### Validation of cross-species comparison of transcriptomic signatures in the POA

We performed snRNA-seq to obtain transcriptomic profiles of POA neurons in hamsters. The POA was dissected from six 10-week-old male Syrian hamsters, and snRNA-seq was conducted using the 10x Chromium platform (Fig. 1a). After quality control, 13,233 nuclei were retained for downstream classification, with a mean of 4979.92 UMI counts, 1978.58 detected genes per cell, and 0.54% mitochondrial transcripts across samples. To enable cross-species transcriptomic comparison, orthologous genes were identified using OrthoFinder2^39^. Of the 21,616 protein-coding genes annotated in the hamster genome, one-to-one orthologs in mice were identified for 16,949 genes (78.4%) (Fig. 1b), whereas 2,036 genes (9.4%) lacked identifiable mouse orthologs, suggesting potential hamster-specific genes (Fig. 1b). To minimize ambiguity in gene correspondence between species, we restricted the cross-species integration to one-to-one orthologous genes, a commonly used strategy in cross-species single-cell analyses^43^, and converted hamster gene symbols to their corresponding mouse ortholog names according to the one-to-one ortholog table.

The resulting dataset was then integrated with published snRNA-seq data from the mouse POA comprising 36,201 cells^28^ using the CCA integration method implemented in Seurat v5^38^. This procedure yielded an integrated gene expression matrix containing 49,434 cells and 14,007 genes (Fig. 1c).

Clustering analysis revealed that major cell classes, including neurons, astrocytes, oligodendrocytes, microglia, and fibroblasts, were clearly distinguished by canonical marker genes (Fig. 1d). Each of these broad cell-type classes contained cells from both species in an intermingled distribution, indicating effective alignment of the major cell classes across species. In addition, the relative proportions of the major cell classes were broadly comparable between the two species (Fig. 1e).

Annotation of excitatory and inhibitory neurons using the established marker genes *Slc17a6* (also known as *vGluT2*) and *glutamate decarboxylase 2* (*Gad2*) reliably distinguished the two neuronal populations (Fig. 1f, g). The relative proportions of these neuronal populations were also highly conserved between species, supporting the robustness of the integrated dataset for subsequent comparative analyses (Fig. 1h). Collectively, these results support broad conservation of the major cellular architecture of the POA between hamsters and mice.

We next examined potential species-specific differences in gene expression profiles. Genes with a fold change > 2 or < 0.5 and *p* < 0.05 were defined as differentially expressed genes between species (species-DEGs). When comparing all cells in the dataset, we identified 321 and 700 species-DEGs enriched in mice and hamsters, respectively. These genes were subjected to gene ontology (GO) enrichment analysis^44,45^, which revealed a significant enrichment of autophagy-related genes among those enriched in hamsters, including *autophagy related 7* (*Atg7*) and *Atg10* (Fig. 1i, j). These results demonstrate that the integrated dataset preserves biologically meaningful species-associated transcriptional signatures, supporting its utility for downstream comparative analyses.

To assess the robustness of neuronal subtype annotation across species, we next subdivided excitatory and inhibitory neurons into 44 and 48 subtypes, respectively, based on discrete marker genes (Fig. 2a, b). This analysis revealed a one-to-one correspondence between species in most excitatory subtypes, except for e21, e22, e30, e33, and e37 (Fig. 2c, left). These mouse-biased subtypes likely reflect sampling bias rather than true species-specific populations, as their spatial distribution suggests localization within the paraventricular thalamic nucleus, the paraventricular hypothalamic nucleus, the bed nucleus of the stria terminalis, and the bed nucleus of the anterior commissure^35^, none of which were included in the hamster samples. Similarly, within inhibitory neurons, while most subtypes were present in both species, five subtypes (i9, i11, i16, i21 and i22) were detected only in the mouse dataset (Fig. 2c, right). These subtypes are localized in the horizontal limb of the diagonal band of Broca or the olfactory tubercle^35^, a region that was not included in our hamster sampling. Apart from these outliers, differences in the relative abundance of subtypes between species were generally modest (Fig. 2c, bottom). Together, these data indicate that mice and hamsters qualitatively share a common set of POA neuronal subtypes, while exhibiting modest species-specific expansion and shrinkage of certain subtypes.

Some species-DEGs were shared across many excitatory or inhibitory subtypes, suggesting common species-enriched genes, whereas others were restricted to only a few subtypes (Fig. 2d). We classified species-DEGs observed in more than half of all subtypes as “shared”, those found in fewer than 10% of subtypes as “cell-type specific,” and the remaining DEGs as “intermediate.” This analysis identified approximately 100 mouse- and hamster-enriched shared species-DEGs, as well as species-DEGs with varying degrees of subtype specificity (Fig. 2d).

We next confirmed the presence of potential thermoregulatory neurons in hamsters that are orthologous to previously characterized neuronal populations in mice. Based on genetic markers, we identified *Adcyap1*- and *Bdnf*-expressing excitatory subtypes in both species, although the expression levels of these genes were generally lower in hamsters (Fig. 3a). Within these subtypes, hamsters showed broader *Lepr* expression and less *Ptger3* expression. Notably, *Opn5* emerged as a conserved marker in both species and distinguished subtype e9 from other excitatory subtypes; this subtype also expressed *Ptger3*. In mice, both *Opn5* and *Ptger3* have been associated with thermoregulation^21,22^. We refer to this subtype as excitatory*^Opn5^*^/*Ptger3*^ neurons. Although *Qrfp* expression was below the detection threshold of snRNA-seq in both species, previous studies have reported substantial overlap among *Qrfp*, *Opn5*, and *Ptger3* expression^23^. Thus, *Opn5* may provide a useful molecular entry for visualizing and manipulating conserved putative thermoregulatory excitatory*^Opn5^*^/*Ptger3*^ neurons in hamsters. The abundance of this subtype did not show a species bias (Fig. 2c). Within excitatory*^Opn5^*^/*Ptger3*^ neurons, we identified both hamster- and mouse-enriched species-DEGs (Fig. 3b). GO analysis revealed enrichment of genes related to GTPase signaling in hamsters and genes associated with synaptic function in mice (Fig. 3c). Consistent with the violin plots (Fig. 3a), the volcano plot showed that *Lepr* expression was higher, whereas *Ptger3* expression was lower in hamsters, implying distinct sensitivities to metabolic state and prostaglandin signaling in hamsters compared with mice (Fig. 3b). In summary, our comparative single-cell transcriptomic analysis highlights the strong conservation of POA cell types in hamsters relative to mice, including populations implicated in thermoregulation that have been characterized in mice.

### Validation of snRNA-seq dataset across non-hibernating and hibernating conditions

To investigate potential changes in gene expression during hibernation, we generated a second cohort including 4–8-month-old male and female hamsters under two housing conditions: (i) a warm condition (12 h light/12 h dark at 21–24 °C), which does not induce hibernation; and (ii) a winter-like condition (8 h light/16 h dark at 4 °C), under which hamsters entered hibernation within approximately two months. We collected two batches for the warm condition (males and females, N = 2 animals per pool) and three batches for the hibernating condition (females, N = 2 animals pooled; males, two batches each containing a single hamster) (Fig. 4a). Of note, during hibernation, hamsters remained curled deep within their nests, exhibited no response to auditory stimuli or cage disturbance, and had markedly reduced body temperature. Samples were collected while the animals were in this state of deep torpor. After quality control, 89,793 nuclei were retained for downstream classification, with a mean of 9950.28 UMI counts, 3375.48 detected genes per cell, and 0.02% mitochondrial transcripts across samples. These datasets were not integrated with the mouse/hamster dataset shown in Fig. 1, as our aim here was to maximize the representation of hamster genes.

We confirmed that both non-hibernating (non-hib) and hibernating (hib) hamsters contained the major cell classes, as defined by canonical marker genes (Fig. 4b, c). Cells from independent biological replicates were well intermixed within each annotated cell population, indicating minimal batch effects across biological replicates. The relative proportions of these major cell classes were highly comparable between the two conditions (Fig. 4d). Similarly, when excitatory and inhibitory neurons were classified as in Fig. 1, their relative proportions remained stable across conditions (Fig. 4e–g). These analyses support the robustness of the seasonal dataset and indicate that the annotation framework established from the cross-species dataset can be consistently applied to an independent hamster cohort.

Condition-associated DEGs (condition-DEGs; fold change > 2 or < 0.5, *p* < 0.05) were subsequently identified to functionally characterize the resulting dataset. At the level of all neurons, 9 upregulated and 259 downregulated condition-DEGs were detected in hibernating hamsters (Fig. 4h). GO analysis revealed that downregulated condition-DEGs were predominantly associated with nucleotide and ribonucleotide metabolic pathways (Fig. 4i). These functional annotations are included as part of the deposited dataset and provide additional characterization of the processed condition-associated gene sets.

We then classified neurons into 50 excitatory and 53 inhibitory subtypes, as in Fig. 2. Among excitatory neurons, the relative proportions of each subtype were largely preserved during hibernation (Fig. 5a, b). Consistent with the global downregulation of gene expression during hibernation (Fig. 4h), downregulated condition-DEGs were more frequently observed using pan-excitatory neuron data (Fig. 5c). The number of condition-DEGs detected in each subtype varied, with only a small fraction of these condition-DEGs shared across >50% of subtypes, whereas most were classified as cell-type specific or intermediate (Fig. 5d). GO analysis revealed that downregulated condition-DEGs in pan-excitatory neurons were associated with nucleotide and ribonucleotide metabolism (Fig. 5e), consistent with the global analysis (Fig. 4i).

Previously described putative thermoregulatory neuronal populations marked by *Adcyap1*, *Bdnf*, *Lepr*, and other genes were also identified in both conditions (Fig. 5f), demonstrating consistent annotation of these neuronal populations within the seasonal dataset. Although their overall abundance was unchanged, the expression levels of *Adcyap1* were significantly downregulated in some excitatory subtypes in hibernating hamsters (Fig. 5f). For example, the volcano plot for subtype 4, corresponding to excitatory*^Opn5^*^/*Ptger3*^ neurons in this dataset, showed an enrichment of downregulated condition-DEGs (Fig. 5g). These genes were also associated with RNA splicing (Fig. 5h), suggesting a potential alteration of gene expression programs in putative thermoregulatory neurons during hibernation.

Similarly, among inhibitory neurons, the relative proportions of each subtype were largely preserved during hibernation (Fig. 6a, b). Condition-DEGs were also predominantly downregulated during hibernation, although certain subtypes, such as inhibitory subtype 15, exhibited abundant upregulated condition-DEGs (Fig. 6c, d). GO analysis revealed enrichment of ribonucleotide metabolism among downregulated condition-DEGs across inhibitory subtypes (Fig. 6e), consistent with the global trend (Fig. 4i). Together, these data provide a transcriptomic landscape of hibernating hamsters relative to non-hibernating animals, establishing a foundation for future functional studies of genes potentially involved in hibernation.

### Validation of the spatial transcriptomics dataset

To characterize the spatial organization of transcriptomic subtypes in the hamster POA, we performed 10x Xenium-based STx under two conditions: (i) a summer-like condition (14 h light/10 h dark at 25 °C), which does not induce hibernation (non-hib); and (ii) a winter-like condition (8 h light/16 h dark at 4 °C), which induces hibernation (hib). We designed a custom panel of 100 Xenium probes targeting representative marker genes of POA neuronal subtypes identified in Fig. 1. In addition, we included a set of IEGs as proxies for neuronal activity, to potentially identify neurons engaged during the induction phase of deep torpor, given that STx approaches can capture state-dependent recruitment using IEG markers such as *c-Fos*^35,46^.

Using this probe set, we conducted Xenium-based STx analysis on POA sections from four 7-month-old female hamsters housed under summer-like conditions and four age-matched animals housed under winter-like conditions undergoing hibernation (Fig. 7a). For the hibernation group, we confirmed that animals had repeatedly entered deep torpor and were sampled approximately 2.5 hours after the onset of body temperature decline following inter-torpor arousal, as monitored by implanted thermal loggers (Fig. 7b). We collected 11–15 sections per animal at 150 μm intervals (Table 1). After quality control and cell segmentation, approximately two million cells were retained for downstream analyses, with a mean of 102.48 transcripts and 21.24 detected genes per cell (Fig. 7a, c).

**Figure 7.**
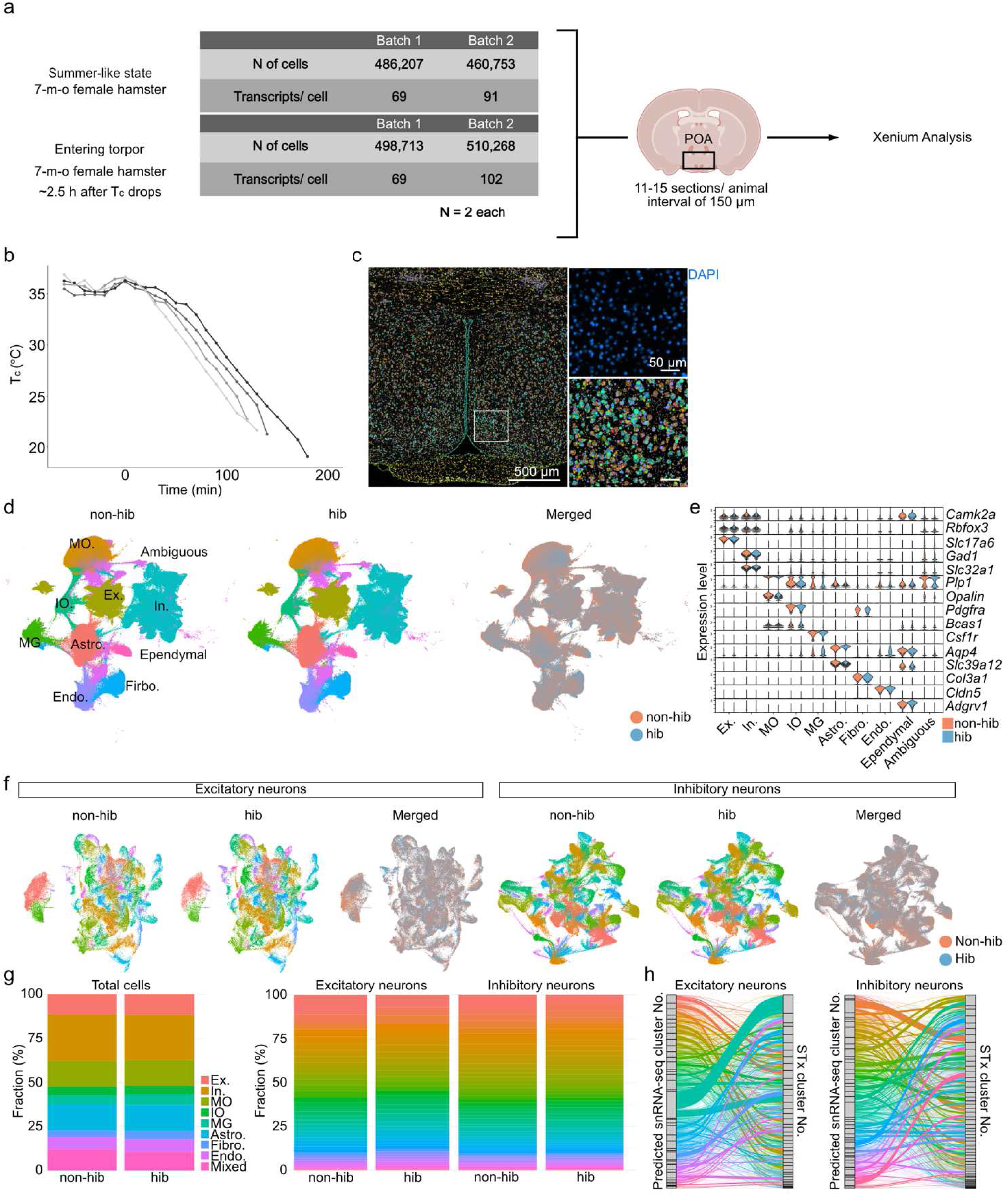
Establishment of an STx atlas of the hamster POA. (**a**) Schematic overview of sample preparation for Xenium-based STx assay. (**b**) Core body temperature (Tc) of hamsters in the hibernation group. The onset of Tc decline toward deep torpor is defined as time 0. (**c**) Fluorescence images of a representative section (left) and a magnified view of the POA (right), stained for DAPI (blue), acquired using the Xenium Analyzer. Colored dots represent transcripts, and colored regions represent cell segments. (**d**) UMAP representation of all cells from non-hibernating hamsters (non-hib, left), hibernating hamsters (hib, middle), and the merged dataset (right). Colors in the left and middle panels indicate cell classes, whereas those in the right panel indicate hibernation state. (**e**) Violin plots showing expression of marker genes for major cell classes. (**f**) UMAP representation of excitatory (left) and inhibitory (right) neurons from non-hibernating hamsters, hibernating hamsters, and the merged dataset. (**g**) Bar graphs showing the proportion of each cell class or neuronal subtype among total cells (left), excitatory neurons (middle), and inhibitory neurons (right). (**h**) Sankey plots showing correspondence between subtypes identified by snRNA-seq and those in the STx dataset, as predicted by RCTD^47^, for excitatory (left) and inhibitory (right) neurons.

**Table 1.**
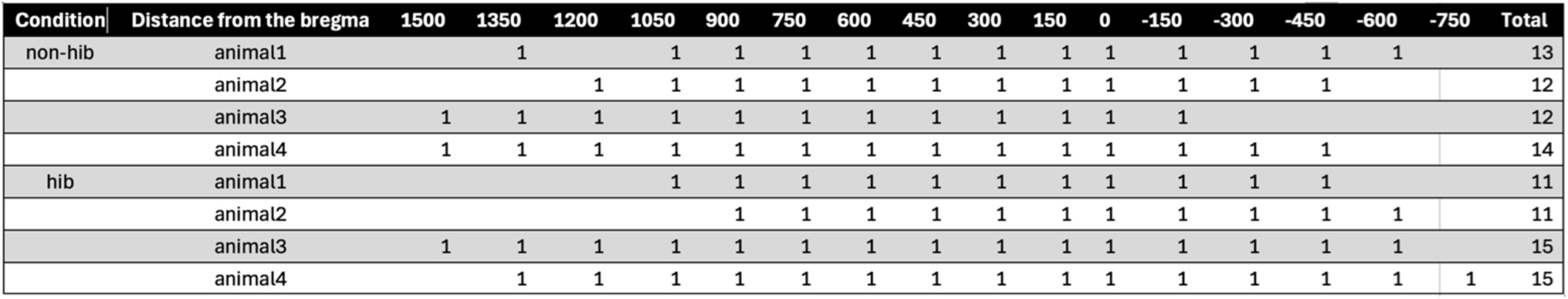
Numbers of POA sections analyzed for the Xenium assay. Rows are organized by condition and animal, and columns list positions ranging from bregma −750 μm to +1500 μm.

The processed STx dataset was integrated using the CCA-based workflow implemented in Seurat v5^38^. Major cell classes were readily identified using canonical marker genes (Fig. 7d, e), and excitatory and inhibitory neurons were classified into 44 excitatory and 57 inhibitory subtypes (Fig. 7f). Cells originating from both experimental conditions were well intermixed within each annotated population, indicating successful integration across biological conditions. The proportions of major cell types and transcriptomic subtypes within excitatory and inhibitory neurons remained stable across conditions (Fig. 7g), consistent with our snRNA-seq data (Figs. 4–6), and cells from both conditions were well intermixed. Furthermore, Robust Cell Type Decomposition (RCTD) analysis^47^ demonstrated strong correspondence between snRNA-seq-defined subtypes (Fig. 2) and STx-derived classifications (Fig. 7h), supporting the robustness of subtype annotation across sequencing modalities.

We next evaluated the spatial distribution of well-characterized thermoregulatory neurons. Among major markers, *Adcyap1*, *Bdnf*, and *Lepr* were broadly expressed in certain excitatory neurons, whereas *Ptger3* and *Opn5* were selectively expressed in only one or two subtypes, most prominently STx subtype E25 (Fig. 8a), corresponding to the excitatory*^Opn5^*^/*Ptger3*^ neurons, consistent with the snRNA-seq results (Figs. 3a and 5f). The STx approach also enabled reliable detection of *Brs3* transcripts, which were most prominently expressed in STx subtype E29, a subpopulation co-expressing *Adcyap1*, *Bdnf*, *Lepr*, and low levels of *Opn5* (Fig. 8a). Spatial mapping of representative thermoregulatory marker genes revealed that excitatory*^Opn5^*^/*Ptger3*^ neurons and *Brs3* neurons were distributed in the hamster anteroventral periventricular nucleus (AVPe) (Fig. 8b), closely resembling the distributions of orthologous populations previously characterized in mice^28,35^. In addition, we observed their distribution in a more posterodorsal region corresponding to part of the median preoptic nucleus (MnPO). In contrast, other subtypes, such as STx subtype E27 (*Adcyap1*+ *Bdnf*+ *Opn5*−), were distributed more ventrally in the medial preoptic nucleus. These data demonstrate that the spatial transcriptomic dataset preserves the anatomical organization of transcriptomic neuronal populations and supports the spatial annotation of previously characterized thermoregulatory cell types.

**Figure 8.**
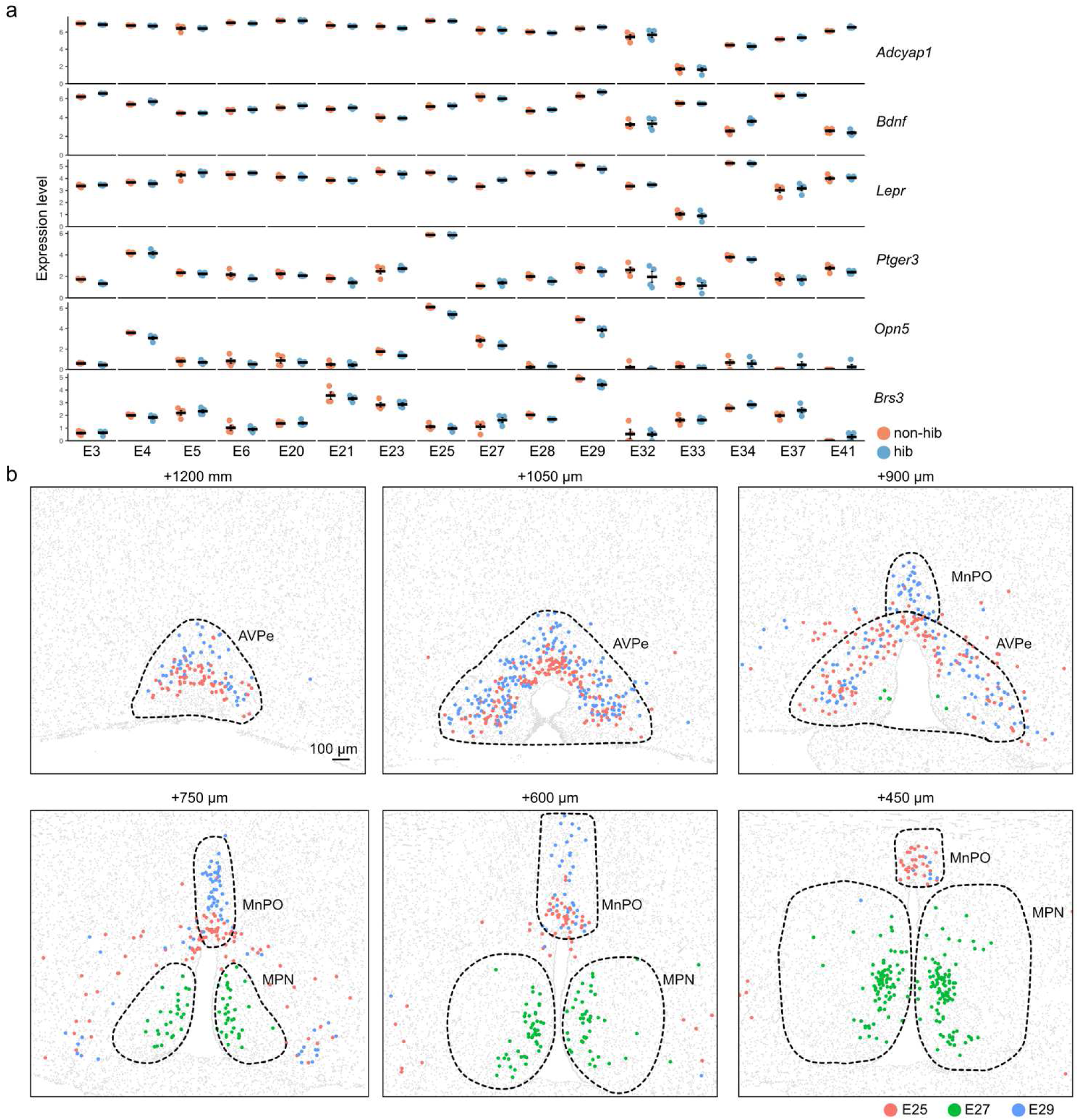
Spatial organization of putative thermoregulation-related neuronal subtypes in the hamster POA. (**a**) Quantification of the normalized expression levels of representative marker genes previously implicated in thermoregulation in mice. Each dot represents the pseudo-bulk expression level from an individual animal (see Methods). Horizontal bars indicate the mean expression level of each marker gene within each subtype, and error bars indicate the SEM. STx excitatory subtype E25, which expresses *Ptger3* and *Opn5*, corresponds to the excitatory*^Opn5^*^/*Ptger3*^ population, whereas subtype E29 shows the strongest expression of *Brs3*. Statistical significance was assessed using the Wilcoxon rank-sum test with Bonferroni correction; no significant differences were detected. N = 4 animals per condition. (**b**) Dot plots showing the representative spatial distributions of STx excitatory subtypes E25, E27, and E29 in the hamster AVPe, MnPO, and medial preoptic nucleus (MPN). Brain-region boundaries (dashed lines) were defined based on the distribution of excitatory subtypes with reference to the hamster brain atlas^52^.

Finally, we evaluated activity-dependent transcription during the early phase of hibernation (2–3 hours after the onset of body temperature decline; Fig. 7b) using two complementary approaches: *c-Fos* expression and an aggregate immediate early gene (IEG) score (see Methods). Under non-hibernating conditions, sparse *c-Fos* expression was detected across subsets of excitatory and inhibitory neuronal populations (Fig. 9a). During early torpor, *c-Fos* expression was globally reduced, and volcano plot analysis further demonstrated significant downregulation of multiple IEGs, including *Arc, c-Fos, Nr4a1, Egr1*, and *Junb* (Fig. 9b). This overall reduction in IEG expression is consistent with previous transcriptomic studies reporting state-dependent suppression of IEGs in the brains of hibernating ground squirrels^48^. Analysis of individual neuronal subtypes likewise showed no subtype with significantly increased *c-Fos* expression, whereas several subtypes exhibited significantly reduced expression during hibernation (Fig. 9c). Consistent with these observations, aggregate IEG scores showed largely reduced or unchanged activity across neuronal subtypes, with only modest increases in the proportion of IEG-positive cells detected in subtypes E5 and I30 (Fig. 9d). Representative spatial distributions of *c-Fos* transcripts together with subtypes E25, E27, and E29 are shown in Fig. 9e. The concordance between *c-Fos* expression and the aggregate IEG score demonstrates consistent activity estimates across complementary analytical approaches, supporting the robustness of activity-dependent transcriptional measurements in the spatial transcriptomic dataset. Collectively, these technical evaluations support the quality and robustness of the spatial transcriptomic dataset and establish a reliable resource for integrative analyses of cell identity, spatial organization, and activity-dependent gene expression in the hamster POA.

**Figure 9.**
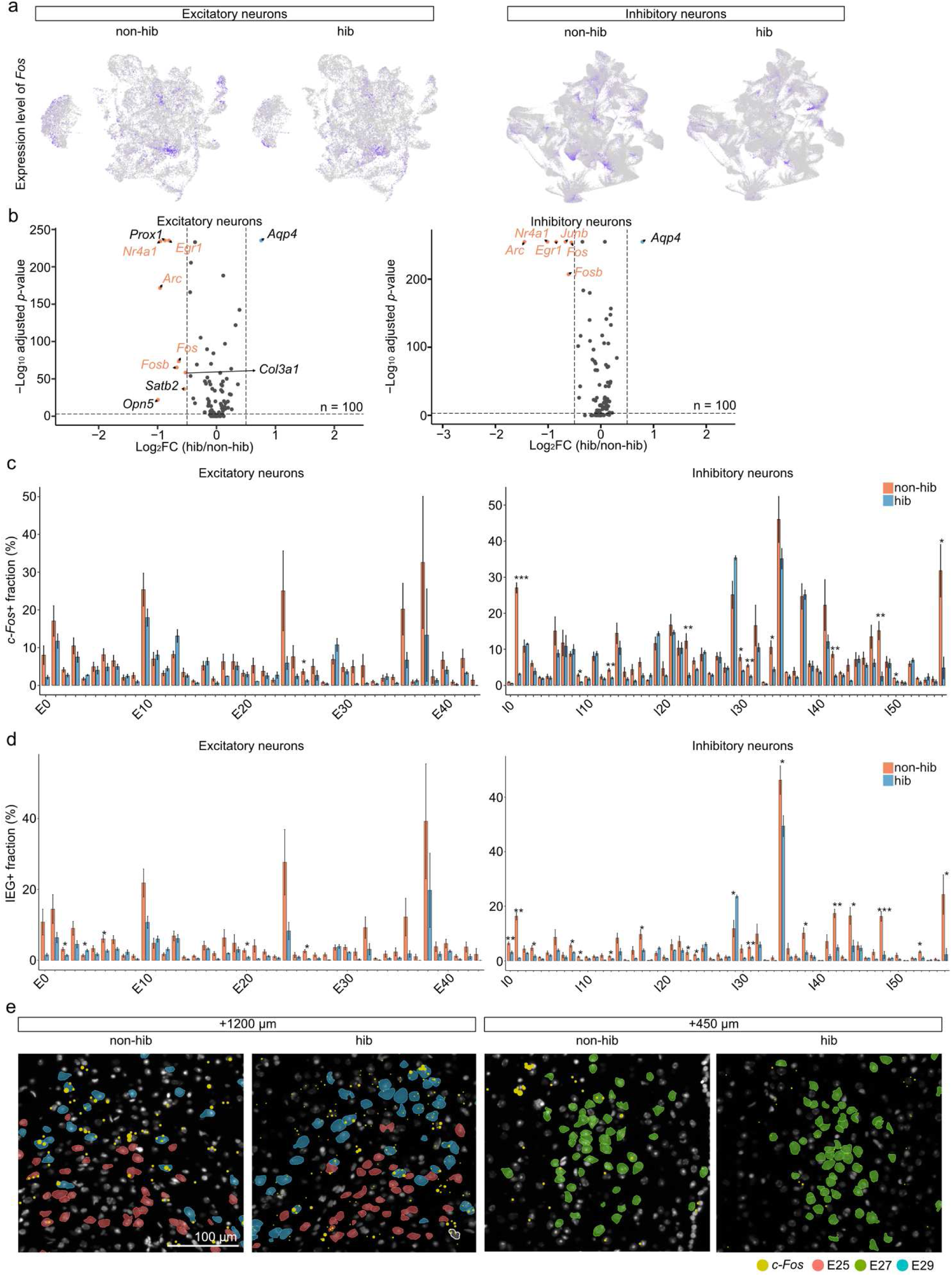
IEG expression in POA neurons during early torpor in hibernating hamsters. (**a**) UMAP representations showing scaled *c-Fos* expression levels in excitatory and inhibitory neurons under each condition (non-hib, non-hibernating; hib, hibernating). (**b**) Volcano plots showing condition-DEGs upregulated (blue dots) and downregulated (orange dots) during hibernation in excitatory (left) and inhibitory (right) neurons. The x-axis shows log_2_-transformed fold change, and the y-axis shows −log_10_-transformed *p*-values from the Wilcoxon rank-sum test. Genes with a fold change > 2 or < 0.5 and *p* < 0.05 were defined as DEGs. IEGs are indicated by orange color. Of note, many IEGs were downregulated in the hibernation group. (**c**, **d**) Bar graphs showing the *c-Fos*+ fraction (**c**) and IEG+ fraction (**d**) in each subtype within excitatory (left) and inhibitory (right) neurons. The *p*-values were calculated using the two-sided Welch’s *t*-test. \**p* < 0.05, \*\**p* < 0.01, \*\*\**p* < 0.001. N = 4 animals per condition. (**e**) Fluorescence images of sections stained for nuclear markers (DAPI; white), acquired using the Xenium Analyzer. Colored regions represent STx excitatory subtypes E25, 27, and 29, with yellow dots indicating *c-Fos* transcripts.

## Usage Notes

One consideration for reuse of the cross-species dataset is that the mouse–hamster comparison in the present study was performed using one-to-one orthologs and CCA-based integration. This pragmatic approach was selected primarily to establish broad correspondence between preoptic cell populations and to facilitate annotation of the hamster dataset. Restricting the analysis to one-to-one orthologs excludes one-to-many and many-to-many homologous relationships and may therefore omit lineage-specific or paralogous transcriptional information. In addition, cross-species integration methods can differ in the extent to which they preserve biological differences while aligning related cell populations^43^. Accordingly, the present cross-species analysis should be interpreted primarily as a framework for cell-type correspondence rather than as an exhaustive analysis of evolutionary divergence between mice and hamsters. More recent approaches that incorporate broader homology relationships^49^ or protein-level representations^50^ may enable finer-resolution comparative analyses of these datasets in the future.

Another limitation of this dataset is that IEG detection by STx did not identify neuronal populations selectively associated with hibernation induction. A recent preprint by Martinez and colleagues^51^ reported an single-cell transcriptome and chromatin-accessibility atlas of the Syrian hamster preoptic area and similarly found broad conservation of preoptic cell types between hamsters and mice. That study further identified preoptic neurons labeled by c-Fos immunohistochemistry during entry into deep torpor and provided functional evidence that these neurons contribute to the hypothermic state. In the present dataset, tissue was collected 2–3 h after the initial decline in Tc, but no robust induction of IEG transcripts was detected by the Xenium assay. Since *Sterile alpha motif domain containing 3* (*Samd3)*-expressing neurons that induce hibernation^51^ were also detected in the current study, this negative result should not be interpreted as evidence that preoptic neurons are inactive during hibernation induction. Several technical and biological factors may account for the discrepancy between transcript-based spatial detection and c-Fos protein labeling. Neuronal activation may be transient and restricted to an early phase of hibernation entry, such that IEG transcript levels had already declined by the time of tissue collection while c-Fos protein remained detectable. Alternatively, neuronal activity may persist, but hypothermia or the associated physiological state may alter the coupling between neuronal activation and IEG transcription. In addition, the sensitivity of the Xenium platform may be insufficient to detect modest IEG induction in this context. Users should therefore exercise caution when using this dataset to infer neuronal activity from IEG transcript abundance. Studies aiming to identify hibernation-associated neuronal activity may benefit from earlier or repeated sampling, protein-based activity mapping, or complementary activity-dependent labeling approaches.

### Quantification and statistics

Statistical tests were performed using custom-written R scripts. All tests were two-tailed. The sample size and statistical tests used are indicated in the figure or figure legends. Error bars are defined in figure legends.

## Data Availability

snRNA-seq and STx data are available in the GEO repository (GSE345405, GSE346081, and GSE345359). All other data generated in this study are presented in the main and Extended data files.

## Code Availability

The custom codes used to analyze the snRNA-seq and STx data in this study are available at Github (https://github.com/gennkenn/snRNA-seq-and-STx-of-the-Syrian-hamster-preoptic-area-across-hibernation-states).

## Acknowledgments

We thank RIKEN R-COMS_BDR for support with the Xenium and snRNA-seq analyses by the Genomics Research and Analysis Support team, Hinako Takase for technical assistance on FFPE preparation and HE staining, and members of the Miyamichi Laboratory for critical reading of the manuscript. H.Sa. is supported by a JSPS postdoctoral research fellowship (PD). This work was supported by the RIKEN BDR-Otsuka Pharmaceutical Collaboration Center (RBOC) founding program to H.Sa. and the JSPS KAKENHI Transformative Research Areas (A) (23H04945, 23H04939) and RIKEN BDR center projects to K.M.

## Author Contributions

H.Sa. and K.M. conceived the experiments. H.Sa. performed the snRNA-seq data acquisition with support from M.T., T.G., and M.H., and conducted the data analysis with support from H.Su. and S.K. H.Sa. also performed STx data collection and analysis with support from D.C., M.K., and T.K. N.S., G.A.S., and H.K. provided hibernating hamsters for snRNA-seq, and C.S., A.Y., and Y.Y. provided hibernating hamster brains for STx.

## Declaration of Interests

The authors declare that they have no competing interests.

## Additional Information

**Correspondence and requests for materials** should be addressed to Kazunari Miyamichi.

## Notes

### Competing Interest Statement

The authors have declared no competing interest.

